# Metabolic engineering of *Escherichia coli* for production of valerenadiene

**DOI:** 10.1101/148338

**Authors:** S. Eric Nybo, Jacqueline Saunders, Sean P. McCormick

**Affiliations:** Ferris State University, College of Pharmacy, Department of Pharmaceutical Sciences, Big Rapids, MI 49307, USA; Ferris State University, Department of Physical Sciences, Big Rapids, MI 49307, USA

**Keywords:** *Valeriana*, valerenic acid, metabolic engineering, sesquiterpene, *Escherichia coli*

## Abstract

*Valeriana officinalis* is a medicinal herb which produces a suite of compounds in its root tissue useful for treatment of anxiety and insomnia. The sesquiterpene components of the root extract, valerenic acid and valerena-1,10-diene, are thought to contribute to most of the observed anxiolytic of Valerian root preparations. However, valerenic acid and its biosynthetic intermediates are only produced in low quantities in the roots of *V. officinalis*. Thus, in this report, *Escherichia coli* was metabolically engineered to produce substantial quantities of valerena-1,10-diene in shake flask fermentations with decane overlay. Expression of the wildtype valerenadiene synthase gene (pZE-*wvds*) resulted in production of 12 μg/mL in LB cultures using endogenous FPP metabolism. Expression of a codon-optimized version of the valerenadiene synthase gene (pZE-*cvds*) resulted in 3-fold higher titers of valerenadiene (32 μg/mL). Co-expression of pZE-*cvds* with an engineered methyl erythritol phosphate (MEP) pathway improved valerenadiene titers 65-fold to 2.09 mg/L valerenadiene. Optimization of the fermentation medium to include glycerol supplementation enhanced yields by another 5.5-fold (11.0 mg/L valerenadiene). The highest production of valerenadiene resulted from engineering the codon-optimized valerenadiene synthase gene under strong P_trc_ and P_T7_ promoters and via co-expression of an exogenous mevalonate (MVA) pathway. These efforts resulted in an *E. coli* production strain that produced 62.0 mg/L valerenadiene (19.4 mg/L/OD_600_ specific productivity). This *E. coli* production platform will serve as the foundation for the synthesis of novel valerenic acid analogues potentially useful for the treatment of anxiety disorders.

## Introduction

*Valeriana officinalis* is a medicinal wild herb indigenous to many habitats, and the root of this plant is used as a nutraceutical preparation (Valerian) that is currently used for the treatment of anxiety and insomnia (Bent et al., 2006). The roots of *V. officinalis* produce a suite of compounds, including valepotriate alkaloids and sesquiterpenes (Bos et al., 1996). Notably, the sesquiterpene components of Valerian root extract are hypothesized to exhibit many of the beneficial anti-anxiety and anti-insomnia effects. Of the sesquiterpenes, valerenic acid is the most potent GABA-A agonist in these extracts, while valerenal, valerenol, and valerena-1,10-diene (valerenadiene) also modulate GABA-A activity to varying extents in zebrafish and mouse models (Del Valle-Mojica and Ortíz, 2012; Takemoto et al., 2014). Most importantly, the anxiolytic effect of valerian has been demonstrated in human clinical studies in recent years (Anderson et al., 2005; Barton et al., 2011).

Valerenic acid in particular has demonstrated nanomolar binding affinity for the GABA-A receptor (Benke et al., 2009). Recently, the putative binding site of valerenic acid has been determined via docking studies and site-directed mutagenesis (Luger et al., 2015). Luger et al. modeled valerenic acid in the GABA-A receptor in a distinct cleft near the β2/3N265 transmembrane residue. The valerenic acid C-12 carboxyl group is predicted to have important hydrogen-bonding interactions with residues β3N265 and β1S265. Furthermore, the C-13, C-14, and C-15 methyl groups of the valerenane skeleton are predicted to have significant hydrophobic interactions within the pocket at residues within this binding pocket (Figure 1) (Luger et al., 2015). In summation, these observations have renewed interest in development of novel valerenic acid analogues for structure activity relationship studies.

**Figure 1.**
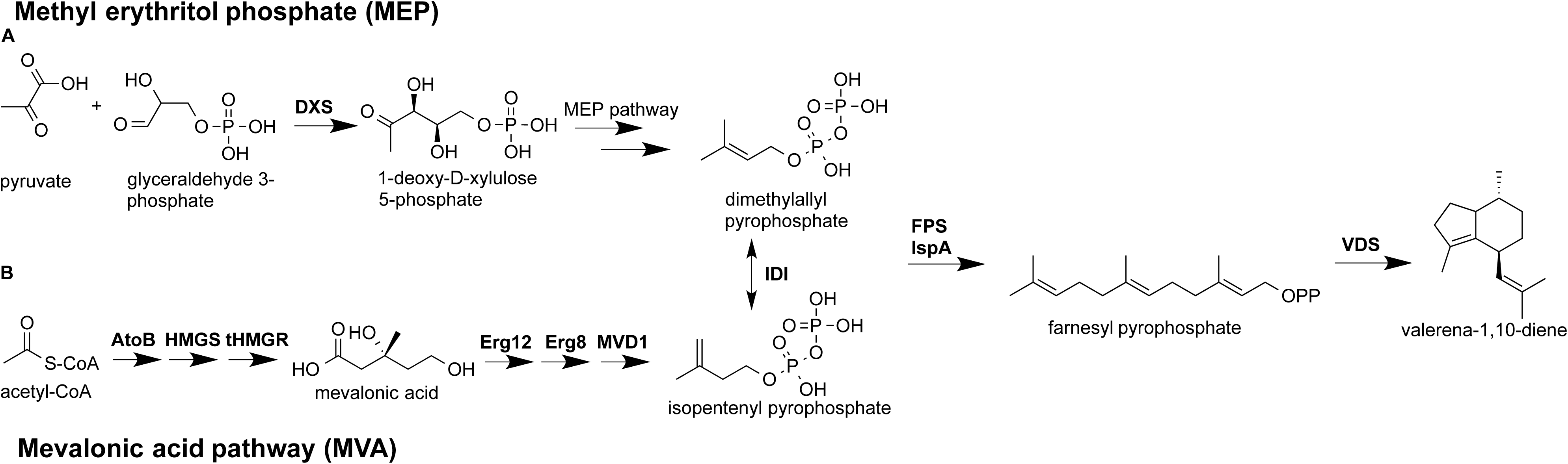
Biosynthesis of valerenadiene via the methyl erythritol phosphate pathway (MEP, pathway A) and the mevalonic acid pathway (MVA, pathway B). IPP and DMAPP are synthesized via the MEP or MVA pathways via different upstream routes. The MEP pathway is initiated via condensation of glyceraldehyde 3-phosphate and pyruvate from glycolysis via 1-deoxy-D-xylulose 5-phosphate synthase (DXS), and the MVA pathway is initiated via condensation of two molecules of acetyl-CoA. Two molecules of IPP and one molecule of DMAPP are condensed into C15 farnesyl pyrophosphate (FPP). FPP is cyclized into valerenadiene via valerenadiene synthase (VDS). The abbreviations of the enzymes are as follows: DXS, 1-deoxy-D-xylulose 5-phosphate synthase; atoB, thiolase; HMGS, hydroxymethylglutaryl-CoA synthase; HMGR, truncated hydroxy-methylglutaryl-CoA reductase; ERG12, mevalonate kinase; ERG8, phosphomevalonate kinase; MVD1, diphosphomevalonate decarboxylase; IDI, isopentenyl diphosphate isomerase; FPS, farnesyl pyrophosphate synthase from *Gallus gallus*; IspA, farnesyl pyrophosphate synthase from *Escherichia coli*; VDS, valerenadiene synthase.

Despite these advances, valerenic acid is produced as a minor constituent in root tissue of *Valeriana officinalis* (0.7-0.9% DW) (Bos et al., 1998). The low production of valerenic acid hinders further attempts at structure activity relationship and biological activity studies. Synthetic routes toward valerenic acid derivatives are also expensive and ecologically unsuitable. Recently, Ricigliano et al. have engineered *V. officinalis* hairy roots for enhanced production of valerenic acid, which lends credence to a metabolic engineering approach for availing these molecules (Ricigliano et al., 2016). Despite this progress, the hairy root system still produces valerenic acid at very lower titers.

As an alternative approach, metabolic engineering of microbial organisms represents a green, cost-effective approach for large-scale production of sesquiterpene pharmaceuticals (Zhang et al., 2011). For example, *Escherichia coli* is one such model host that affords several advantages over plant-based systems due to its fast growth kinetics and capacity to produce high-value chemicals via fermentation on simple carbon sources (Lee, 1996). Furthermore, *E. coli* boasts considerable genetic tools, including multiple promoters and expression vectors that establish it as an ideal host for metabolic engineering. Additionally, terpenes are attractive molecules for metabolic engineering, due to their use as fragrances, flavors, and advanced biofuels (Peralta-Yahya et al., 2011; Sowden et al., 2005). Subsequently, these observations established *E. coli* as a suitable platform for engineering of valerenic acid in this present study.

Two specialized isoprenoid biosynthetic pathways exist for production of C5 isoprenyl phosphate precursors, dimethyl allyl pyrophosphate (DMAPP) and isopentenyl pyrophosphate (IPP). Many bacteria, algae, and plant chloroplasts employ the methyl erythritol phosphate pathway (MEP), which fluxes glyceraldehyde-3-phosphate and pyruvate towards IPP and DMAPP (Figure 1) (Rohmer, 1999). Fungi and non-plant eukaryotes utilize the mevalonate pathway (MVA) to convert acetyl-coenzyme A (acetyl-CoA) to IPP and DMAPP via eight enzymatic steps (Figure 1) (Martin et al., 2003). Subsequently, IPP and DMAPP are concatenated into progressively longer C10, C15, or C20 molecules by prenyltransferase enzymes. For sesquiterpene metabolism, farnesyl pyrophosphate synthase condenses 2 IPP units and 1 DMAPP unit to produce farnesyl pyrophosphate (FPP). Subsequently, FPP can be cyclized by a variety of sesquiterpene hydrocarbons by cognate terpene synthases, such as valerenadiene synthase (VDS) (Figure 1).

However, *E. coli* generates only a finite pool of FPP, and while introduction of a sesquiterpene synthase results in detectable production of sesquiterpenes (Martin et al., 2001), overproduction of the molecule requires redirecting substantial carbon flux to the limited substrate FPP. For example, amorphadiene is a sesquiterpene intermediate in the biosynthesis of the antimalarial drug artemisinin. Martin and co-workers discovered that heterologous expression of the mevalonate isoprenoid pathway from yeast in *E. coli* leads to unregulated carbon flux towards FPP and amorphadiene, which can be semi-synthetically converted to artemisinin (Martin et al., 2003). In using this substrate-engineering approach, Keasling and co-workers have produced amorphadiene in *E. coli* at yields of 500 mg L^-1^ in shake flask (Redding-Johanson et al., 2011) and 27 g L^-1^ in bioreactors (Tsuruta et al., 2009). This generalized approach lends itself to the microbial synthesis of other sesquiterpenes via introduction of variant terpene synthases.

In this report, a metabolic engineering platform was developed for synthesis of valerenadiene in *E. coli* in three steps. Because plant terpene synthase genes are poorly expressed in *E. coli*, owing to different codon usage, first an *E. coli* codon-optimized version of the valerenadiene synthase gene (*cvds*) was synthesized. This lead to a three-fold higher production of valerenadiene over expression of the wildtype plant gene (*wvds*). Secondly, we enhanced carbon flux to FPP by designing a construct that overexpressed several rate-limiting steps of the native methyl erythritol phosphate (MEP) pathway. Thirdly, we cloned *cvds* under the control of P_trc_ and P_T7_ promoters to further enhance terpene synthase protein levels and we co-expressed these constructs with a heterologous mevalonate pathway from baker’s yeast. These efforts resulted in a high-level production strain that synthesized 62.0 mg L^-1^ valerenadiene (19.4 mg/L/OD_600_ specific productivity) in shake flask cultures.

## Methods and Materials

### Bacterial strains and growth conditions

*E. coli* JM109 (New England Biolabs) was used as the host for all routine cloning manipulations, and *E. coli* DH5αZ1 (Expressys, Germany) and *E. coli* BL21(DE3) (ThermoFisher) were used as hosts for sesquiterpene production (Supplementary Table 1). *E. coli* DH5αZ1 overexpresses a copy of the lacIq repressor on the chromosome for efficient repression of the lac operator (Lutz and Bujard, 1997). Chemically competent *E. coli* were generated with the *E. coli* Mix and Go Transformation Kit (ZYMO Research) and were transformed using standard molecular methodologies (Sambrook and W Russell, 2001). *E. coli* strains were grown in LB agar or LB broth at 37 C for routine maintenance. For production of sesquiterpenes, *E. coli* DH5αZ1 derivatives were grown in 2xYT with supplemented glycerol at 30 C. for production of valerenadiene. For expression of the mevalonic acid pathway, bacterial growth media was buffered with phosphate buffered saline using a stock solution of 10x PBS (Sambrook and W Russell, 2001). Strains were supplemented with ampicillin (100 μg mL^-1^), chloramphenicol (35 μg mL^-1^), and kanamycin (50 μg mL^-1^) as necessary. When multiple plasmids were co-expressed, the chloramphenicol and kanamycin concentrations were adjusted to one-half these amounts.

### Cloning of vds and engineered MEP pathway constructs

Oligonucleotide primers were synthesized by IDT-DNA (Supplementary Table 2), and sequences were verified by sequencing analysis (ACGT, Inc.). Polymerase chain reaction was carried out using Primestar^®^ HS Polymerase (Takara Bio.) by following the manufacturer’s protocols. The wildtype valerenadiene synthase gene (*wvds*) gene was amplified via polymerase chain reaction from the pET28a(+)-VoTPS1 (hereafter referred to as pET-*wvds*) construct as described previously (Yeo et al., 2013). The *cvds* gene was codon-optimized for expression in *E. coli*, synthesized, and spliced into cloning vector pUC57-*cvds* (GenScript). The *cvds* gene was amplified via polymerase chain reaction, and both the *wvds* and *cvds* genes were cut and spliced into the *Eco*RI/*Bam*HI restriction sites of pZE12MCS under the control of the intermediate strength P_LlacO1_ promoter (Expressys, Germany) to afford constructs pZE-wVDS and pZE-cVDS, respectively (Lutz and Bujard, 1997). For insertion under the stronger P_trc_ promoter, *cvds* was PCR amplified, cut, and spliced into the *Bam*HI/*Eco*RI sites of pTrcHisA (Thermo Fisher Scientific) to afford pTrcHis-*cvds*. In this construct, *cvds* is fused to the *N*-terminal hexahistidine and Xpress™ epitope tags. For comparisons involving the PT7 promoter, the *cvds* gene was cloned into the *Nco*I and *Eco*RI sites of pET28a(+) to afford pET-*cvds*. pET-*wvds* features *wvds* fused to the PT7 promoter and was included in experiments to reflect baseline production of valerenadiene from T7-polymerase driven expression of the wildtype gene.

The *dxs*-*idi*-*fps* genes were synthesized as a polycistron as previously described in vector pUC57-operon 3 (Genscript) (Bell et al., n.d.). The *dxs-idi-fps* genes were digested with *Eco*RI/*Bam*HI and were spliced into the *Eco*RI/*Bam*HI sites of pSTV28 to afford pSTV-*dxs-idi-fps*.

### Cloning and expression of mevalonate pathways

pBbA5c-MevT(CO)-MBIS(CO, ispA) was a gift from Jay Keasling & Taek Soon Lee (Addgene plasmid # 35151). For co-expression of valerenadiene synthase and the mevalonate pathway, *E. coli* competent cells were co-transformed with the relevant constructs and plated on LB agar supplemented with antibiotics. For terminal endpoint assays, *E. coli* strains were grown in 1 mL of LB supplemented with antibiotics for 12-16 hours until the strains were in stationary phase growth, and then they were inoculated into 5 mL 2x YT medium containing 3% glycerol with an overlay of 1 mL of decane. For time course assays, *E. coli* strains were grown in 50 mL 2x YT medium containing 3% glycerol with an overlay of 10 mL of decane. Time points were collected at 12, 24, 36, 48, 72, and 96 hours of 100 microliters of culture broth (for OD_600_ measurements) and 100 microliters of decane overlay for valerenadiene determination.

### GC-MS analysis of valerenadiene

Valerenadiene production by the various strains was measured by GC-MS via selective ion monitoring for the molecular ion (204 *m/z*), the 189 *m/z* ion, and the 69 *m/z* ion, as described previously (Martin et al., 2003). Cells were grown in LB or 2x YT medium with 3% glycerol and induced with 200 micromolar IPTG to express the *wvds* or *cvds* gene and either the engineered MEP operon or the mevalonate pathways. Cultures were covered with a 20% (v/v) decane overlay to trap volatile sesquiterpenes. 100 microliters of decane overlay was diluted in 900 microliters of hexane, and 1 microliter of sample was subjected to GC-MS analysis. Samples were analyzed at the Ferris State University Shimadzu Core Laboratory for Academic Research Excellence (FSU-SCLARE) on a Shimadzu QP-2010 Ultra GC mass spectrometer. Valerenadiene in experimental samples was compared to authentic valerenadiene standard. Valerenadiene concentration was converted to beta caryophyllene equivalents using a caryophyllene standard curve and the relative abundance of 69 *m/*z, 189 *m/z*, and 204 *m/z* mass fragments. Valerenadiene standard was isolated as described previously (Yeo et al., 2013).

## Results

### Valerenadiene production in E. coli and gene optimization

To evaluate the production of valerenadiene synthase in *E. coli*, the gene encoding wildtype valerenadiene synthase (*wvds*) from *V. officinalis* was cloned under the control of the IPTG-inducible P_LlacO1_ promoter of high copy number expression plasmid pZE12MCS to afford pZE-wVDS. Previously, other groups have reported that sesquiterpene production in *E. coli* is limited by poor expression of cognate plant terpene synthases (Martin et al., 2001; Wang et al., 2011). We surmised that the *wvds* gene may encode several rare codons that inhibit optimal translation in *E. coli*. Therefore, we synthesized a codon-optimized version of the valerenadiene synthase gene for expression in *E. coli* (GenScript) and fused it to the P_LlacO1_ to afford pZE-cVDS. pZE-wVDS, pZE-cVDS, and empty pZE12MCS vector were transformed in *E. coli*. The *E. coli* lines were grown in 5 mL of LB medium overlaid with decane for 48 hours and induced with IPTG. A major valerenadiene peak was detected in extracts of pZE-wVDS and pZE-cVDS cultures that eluted at the same retention time and exhibited identical mass fragmentation pattern to valerenadiene standard (Supplementary Figure 1). This peak was not detected in pZE12MCS control strains, which confirmed low production of valerenadiene in the VDS-engineered lines. *E. coli* pZE-wVDS produced 8 μg/L and 12 μg/L valerenadiene at 24 and 48 hours, respectively, whereas *E. coli* pZE-cVDS produced 16 and 32 μg/L valerenadiene at 24 and 48 hours, respectively (Figure 2). This results suggests that the codon-optimized valerenadiene synthase is more efficiently expressed than the wildtype version, which results in a 3-fold improvement in valerenadiene titer.

**Figure 2.**
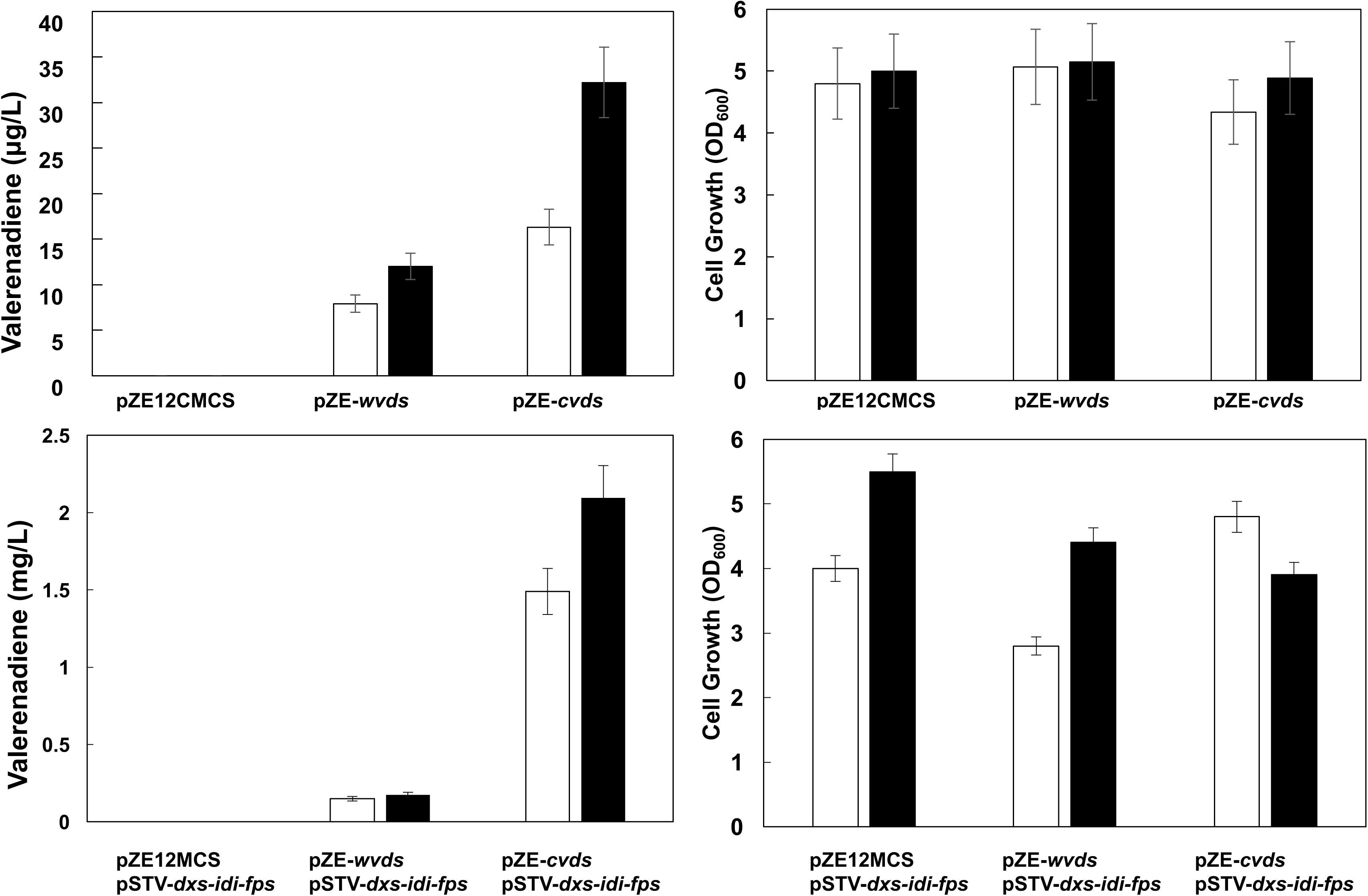
Engineering of wildtype (pZE-*wvds*) and codon-optimized (pZE-*cvds*) valerenadiene synthase. **(A)** *E. coli* engineered with empty vector control, wildtype, and codon optimized VDS genes was grown for 24 (white bars) and 48 hours (black bars) and yield of valerenadiene was determined. **(B)** Growth of engineered *E. coli* lines was determined by measuring optical density (OD_600_) at 24 (white bars) and 48 hours (black bars), respectively. **(C)** *E. coli* strains were co-transformed with empty vector control, wildtype, or codon-optimized *vds* and pSTV-*dxs-idi-fps* for enhancement of the endogenous MEP pathway. (**D**) Growth of engineered *E. coli* lines was determined by measuring optical density (OD_600_). Strains were inoculated in 5 mL cultures of LB media with 1 mL decane overlay and experiments were carried out in triplicate. Error bars indicate standard error of the mean.

However, the yields of valerenadiene achieved were quite low, and we surmised this was likely due to low levels of prenyl phosphate precursors (IPP, DMAPP, and FPP). The endogenous MEP pathway of *E. coli* produces a small pool of FPP that is the precursor for *trans*-octaprenyl diphosphate (ODP) in production of ubiquinone and *cis, trans*-undecaprenyl diphosphate (UDP) for production of peptidoglycan (Asai et al 1994, Bouhss et al 2008, Okada et al 1997). However, the production of FPP by the endogenous pathway is limited due to regulation of several key steps (Estévez et al., 2001). Therefore, we designed a construct that would overexpress rate-limiting steps of the MEP pathway to augment greater carbon flux from primary metabolism to FPP. In our previous work, we engineered *Rhodobacter capsulatus* to produce high levels of the triterpene botryococcene, which is biosynthesized from two molecules of FPP via squalene synthase-like enzymes (SSL-1 and SSL-3), via expression of the deoxy-xylulose phosphate synthase (*dxs*), isopentenyl diphosphate isomerase (*idi*) genes from *E. coli*, and farnesyl pyrophosphate synthase (*fps*) gene from *Gallus gallus* (Khan et al., 2015). Fusion of the *dxs*-*idi*-*fps* genes to the SSL-1+3 chimeric botryococcene synthase and expression of the resulting construct generated 5 mg/gDW production of botryococcene in the *R. capsulatus* host. In a similar fashion, we fused the *dxs*-*idi*-*fps* genes under the control of the IPTG-inducible P_lacUV5_ promoter of pSTV28 to generate construct pSTV-*dxs-idi-fps*. This construct was co-transformed with pZE-*wvds* and pZE-*cvds*, and the lines were grown in 5 mL LB medium with decane overlay at 30°C for 48 hours and induced with IPTG. Co-expression of pSTV-*dxs*-*idi*-*fps* with pZE-wVDS and pZE-cVDS resulted in a 14- fold increase (0.17 mg/L) and a 65-fold increase (2.09 mg/L) in production of valerenadiene, respectively (Figure 2). These results demonstrate that expression of the engineered MEP pathway greatly augmented carbon flux to FPP and valerenadiene.

### Effect of glycerol supplementation on valerenadiene production

With the engineered MEP pathway construct in hand, we hypothesized that optimization of the production medium could further enhance sesquiterpene yield. In fact, Zhang et al. reported that glycerol supplementation of culture medium increased sabinene production to >40 mg/L in *E. coli* engineered with sabinene synthase and a heterologous mevalonate pathway (Zhang et al., 2014). Furthermore, Morrone and co-workers engineered higher levels of _C20_ abietadiene production and increased biomass when glycerol supplemented media was used (e.g. increasing abietadiene specific productivity from 1 mg/L/OD_600_ without glycerol supplementation to 2.5 mg/L/OD_600_ with glycerol supplementation) (Morrone et al., 2010). *E. coli* pZE-wVDS/pSTV-*dxs-idi-fps* and *E. coli* pZE-cVDS/pSTV-*dxs-idi-fps* were grown in 5 mL of 2x YT medium supplemented with 0 to 5% glycerol. The *E. coli* pZE-wVDS/pSTV-*dxs-idi-fps* valerenadiene production increased 15-fold from 0.11 mg/L to 1.5 mg/L when supplemented with glycerol, however, further production was possibly stymied by poor heterologous expression of valerenadiene synthase. In contrast, *E. coli* pZE-cVDS/pSTV-*dxs-idi-fps* demonstrated better responsiveness to the glycerol supplementation, as valerenadiene productivity increased from 5.1 mg/L to 11.0 mg/L when supplemented with 3% glycerol (Figure 3). Therefore, 3% glycerol was used for all further experiments.

**Figure 3.**
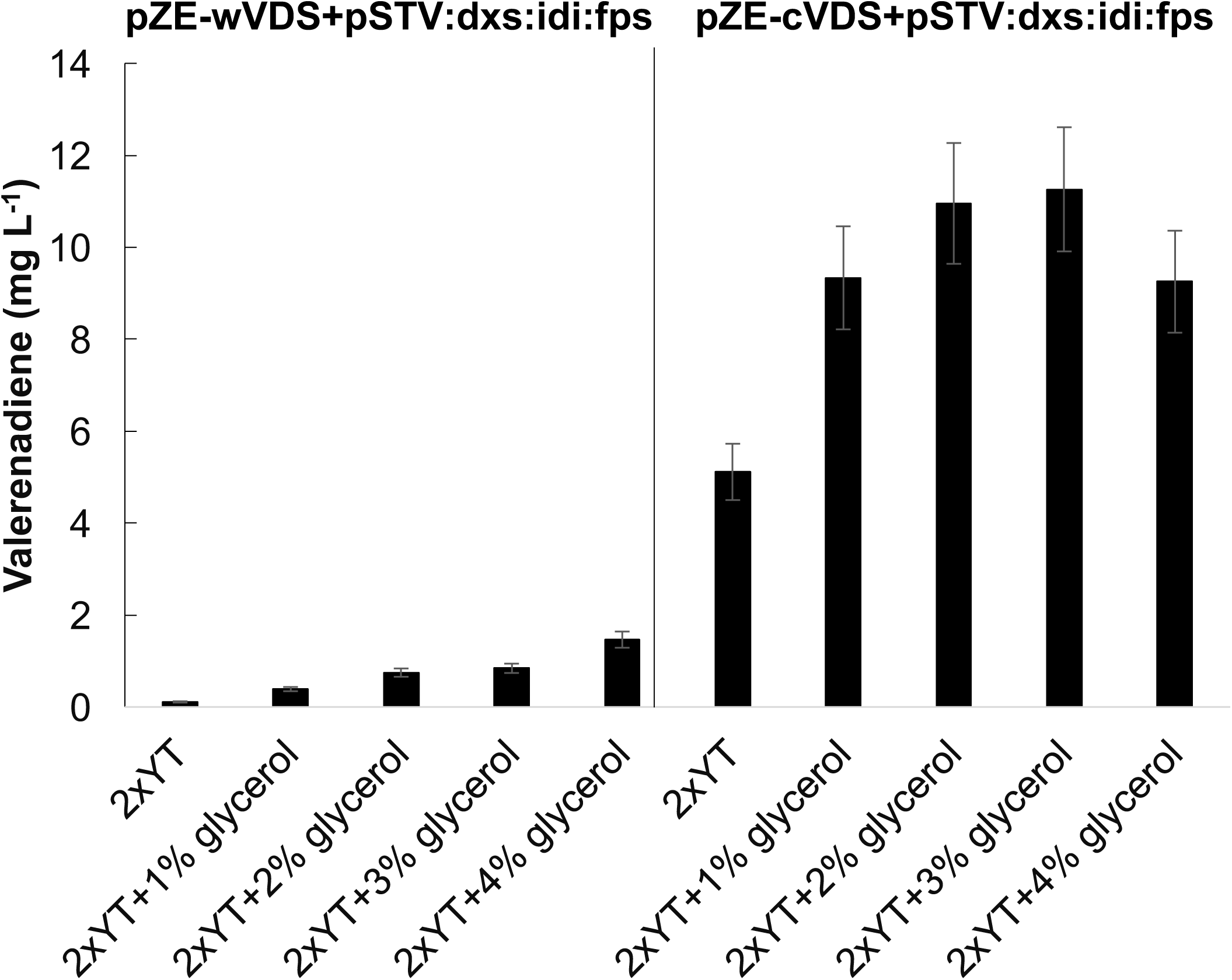
Effect of glycerol conditions on production of valerenadiene. *E. coli* strains engineered with pZE-*wvds* and pSTV-*dxs-idi-fps* or pZE-*cvds* and pSTV-*dxs-idi-fps* were grown in 5 mL 2xYT media with 0%-4% glycerol and 1 mL decane overlay for 48 hours and valerenadiene yield was determined. Experiments were carried out in triplicate and error bars indicate standard error of the mean.

To observe production of valerenadiene over time, we conducted a time course of *E. coli* pZE-wVDS/pSTV-*dxs-idi-fps* and *E. coli* pZE-cVDS/pSTV-*dxs-idi-fps* in 2x YT+3% glycerol media over 72 hours. The results of the time course replicated the observations in the glycerol supplementation experiment. *E. coli* co-expressing the engineered MEP pathway and the *wvds* gene reached maximal growth (OD_600_=9.0) and valerenadiene production (3 mg/L) at 24 hours (Figure 4). The *E. coli* line co-expressing the engineered MEP pathway and *cvds* gene reached also reached both its greatest level of growth (OD_600_ =9.0) and valerenadiene production (10.6 mg/L) at 24 hours. Valerenadiene concentration decreased in both lines from 48 to 72 hours, likely due to volatilization of the sesquiterpene.

**Figure 4.**
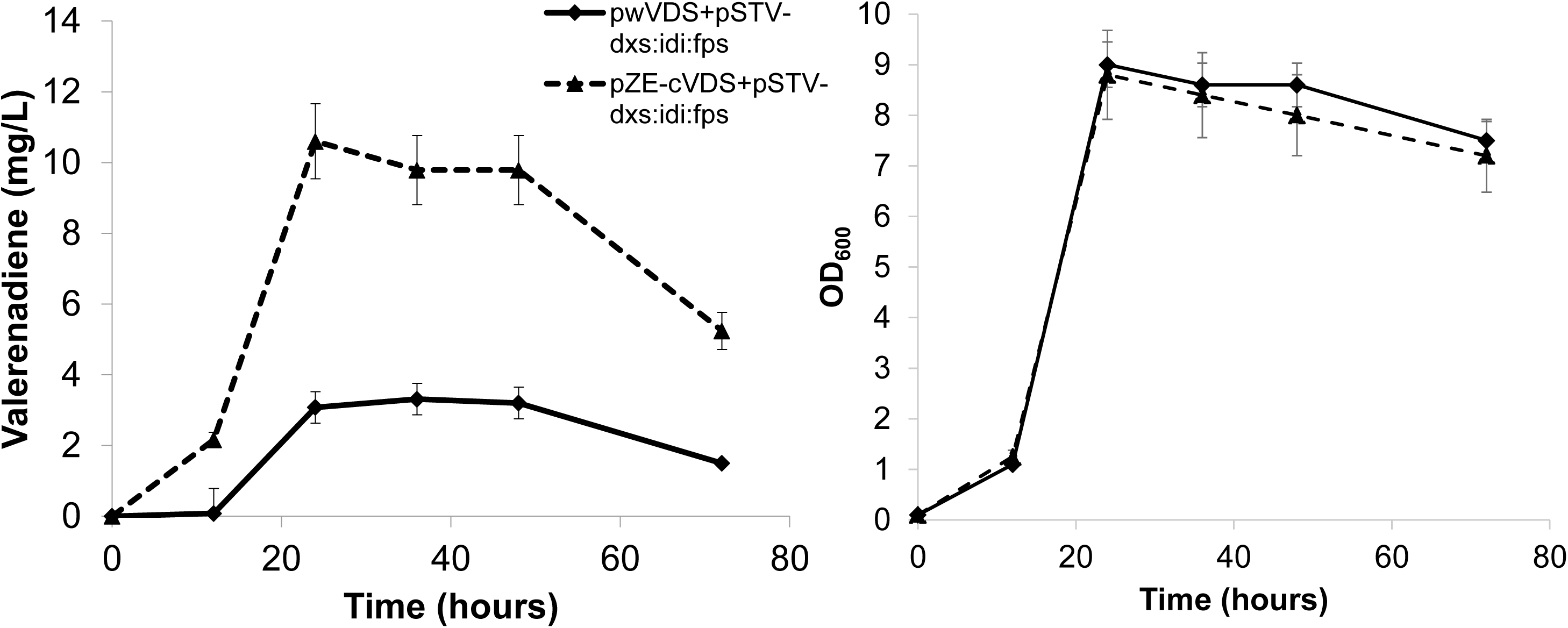
Time course of *E. coli* expressing an engineered MEP pathway and valerenadiene synthase. *E. coli* DH5αZ1 strains harboring the engineered MEP construct pSTV-*dxs-idi-fps* were co-expressed with the wildtype (pZE*-wvds,* solid line) or codon-optimized (pZE-*cvds*, dotted line) valerenadiene synthase gene. Cultures were grown in 50 mL 2x YT+3% glycerol with 10 mL decane overlay for 72 hours. (Left panel) Valerenadiene yield (mg L^-1^) and (right panel) biomass (OD_600_) were determined from double triplicate experiments. Error bars indicate the standard error of the mean.

### Engineering of vds under the P_trc_ promoter

We hypothesized that low-level expression of the terpene synthase from the P_LlacO1_ promoter might be a metabolic rate-limiting factor in the turnover of FPP substrate. Based on the previous experiment, expression of the *cvds* gene resulted in 5-10-fold higher quantities of valerenadiene than expression of *wvds*, depending on glycerol supplementation, which was inferred to be the direct result of higher concentration of terpene synthase catalyst in the *E. coli* cell. Therefore, the *cvds* gene was fused to the strong P_trc_ promoter to afford construct pTrcHis-*cvds*. This construct was expressed alone and co-expressed with pSTV-*dxs-idi-fps* in *E. coli* DH5αZ1 cells in triplicate 5 mL 2xYT+3% glycerol fermentations for 48 hours. The strain harboring pTrcHis-*cvds* produced 0.046 mg/L valerenadiene, whereas the strain harboring pTrcHis-*cvds* and pSTV-*dxs-idi-fps* produced approximately 20.3 mg/L (Supplementary Figure 2). This represents a 440-fold improvement over the pTrcHis-*cvds* only expression line. Furthermore, expression of the *cvds* gene by the P_trc_ promoter (e.g. by strain *E. coli* DH5αZ1/(pTrcHis-*cvds*)/(pSTV-*dxs-idi-fps*)) represented a 40% increase in valerenadiene titers as compared to expression from P_LlacO1_ (e.g. by strain *E. coli* DHαZ1 /(pZE-*cvds*)/(pSTV-*dxs-idi-fps*)).

Recently, the MEP pathway has been demonstrated to be subject to endogenous regulation by *E. coli*, possibly due to feedback inhibition of the metabolite methylerythritol cyclodiphosphate (MEcDP) (Banerjee and Sharkey, 2014). We rationalized that FPP enhancement might be limited via the MEP pathway, therefore we co-expressed an exogenous MVA pathway to greatly augment FPP precursor levels along with pTrcHis-*cvds*. In a previous study, Peralta-Yahya and co-workers engineered a heterologous MVA pathway for production of the sesquiterpene biofuel bisabolene (Peralta-Yahya et al., 2011). Expression of the construct pBbA5c-MevT(CO)-MBIS(CO, ispA) along with the codon-optimized AgBis gene resulted in production of 586±65 mg/L bisabolene. pBbA5c-MevT(CO)-MBIS(CO, ispA) features a codon-optimized, eight gene MVA pathway driven by the P_trc_ promoter on a medium copy number plasmid (Supplementary Table 1). The three upstream gene products (*atoB*, *hmgs*, *thmgr*) convert acetyl-CoA to mevalonic acid, and the five downstream gene products (*erg12*, *erg8*, *mvd1*, *idi*, *ispA*) convert mevalonic acid to farnesyl pyrophosphate (Figure 1). Similarly, *E. coli* DH5αZ1 was transformed with pTrcHis-*cvds* and pBbA5c-MevT(CO)-MBIS(CO, ispA) and grown in triplicate 5 mL 2xYT+3% glycerol fermentations for 48 hours. This strain produced 42.5 mg/L valerenadiene, a 1000-fold increase over the *cvds*-expressing line and a 2-fold increase over the engineered MEP pathway (Supplementary Figure 2). This result demonstrated that the mevalonic acid pathway efficiently coupled the microbial synthesis of FPP to valerenadiene synthase to produce significant levels of valerenadiene. Thus, the mevalonic acid pathway was used for further experiments. These results were further confirmed in a time course experiment of the same strain, which achieved a titer of 35 mg/L valerenadiene at 24 hours (Figure 5).

**Figure 5.**
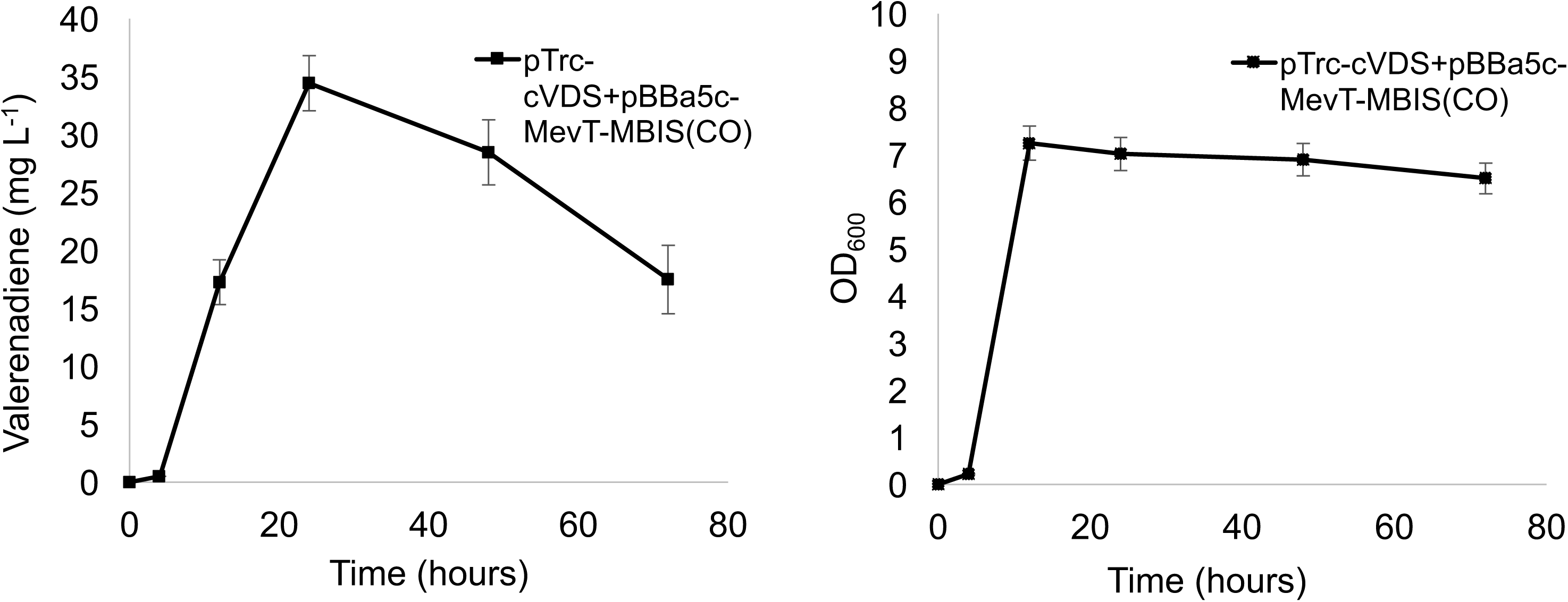
Time course experiment of *E. coli* co-expressing pTrcHis-*cvds* and the *Saccharomyces cerevisiae* MVA pathway. *E. coli* DH5ΑZ1 strains harboring the pBBa5c-MevT+MBIS(CO,IspA) and pTrcHis-*cvds* constructs were grown in 50 mL 2x YT+3% glycerol with 10 mL decane overlay for 72 hours. (Left panel) Valerenadiene yield (mg L^-1^) and (right panel) biomass (OD_600_) were determined from double triplicate experiments. Error bars indicate the standard error of the mean.

### Co-expression of vds with the T7 RNA polymerase promoter and MVA pathway

We observed that the P_trc_-*cvds* gene constructs demonstrated more robust production of valerenadiene than the comparatively weaker P_LacO1_-*cvds* gene constructs, and we inferred that levels of VDS catalyst may be rate-limiting in this latter system. Therefore, we decided to compare the expression of the *wvds* and *cvds* genes from a “weak” promoter system (P_LacO1_), an “intermediate/strong” promoter system (P_trc_), and a strong promoter system (P_T7_) along with the mevalonic acid pathway. We hypothesized that the P_LacO1_ promoter might not be able to drive sufficient expression of VDS to efficiently turnover the abundant FPP substrate synthesized via the mevalonic acid pathway. *E. coli* DH5αZ1 pZE-*wvds* and *E. coli* DH5αZ1 pZE-*cvds* were grown in triplicate cultures as before and produced minimal amounts of valerenadiene (Supplementary Figure 3. When pBbA5c-MevT(CO)-MBIS(CO, ispA) was co-expressed in these lines, they produced 2.8 mg/L (0.497 mg/L/OD_600_) and 7.3 mg/L (1.03 mg/L/OD_600_), respectively, which strongly suggested that the amount of VDS catalyst was limited (Supplementary Figure 3). Expression of *cvds* from the P_trc_ promoter with the mevalonic acid pathway increased production 6-fold to 42.5 mg/L (6.07 mg/L/OD_600_) (Supplementary Figure 3).

Subsequently, the *wvds* and *cvds* genes were cloned into pET28a for expression under the strong T7 RNA polymerase promoter. The resulting constructs, pET-*wvds* and pET-*cvds* were introduced into *E. coli* BL21(DE3) cells to utilize the IPTG-inducible T7 RNA polymerase system. These plasmids were transformed alone or in combination with the mevalonic acid pathway and grown in triplicate 2xYT + 3% glycerol cultures for 48 hours. The pET-*wvds*-only line produced 0.345 mg/L valerenadiene, whereas the pET-*cvds*-only line produced 1.09 mg/L valerenadiene (Figure 6). Co-expression of pET-*wvds* and pBbA5c-MevT(CO)-MBIS(CO, ispA) resulted in production of 10.5 mg/L valerenadiene (3.27 mg/L/OD_600_) (Figure 7). Most importantly, when pET-*cvds* was introduced with the mevalonic acid pathway, the highest-yielding valerenadiene strain was achieved with production of 62.0 mg/L valerenadiene (19.4 mg/L/OD_600_ specific productivity). These experiments demonstrated that the amount of VDS catalyst and FPP substrate *in vivo* are both limiting factors for production of valerenadiene, which can be remedied via expression of *cvds* from a strong P_trc_/_PT7_ promoter and heterologous expression of the mevalonic acid pathway.

**Figure 6.**
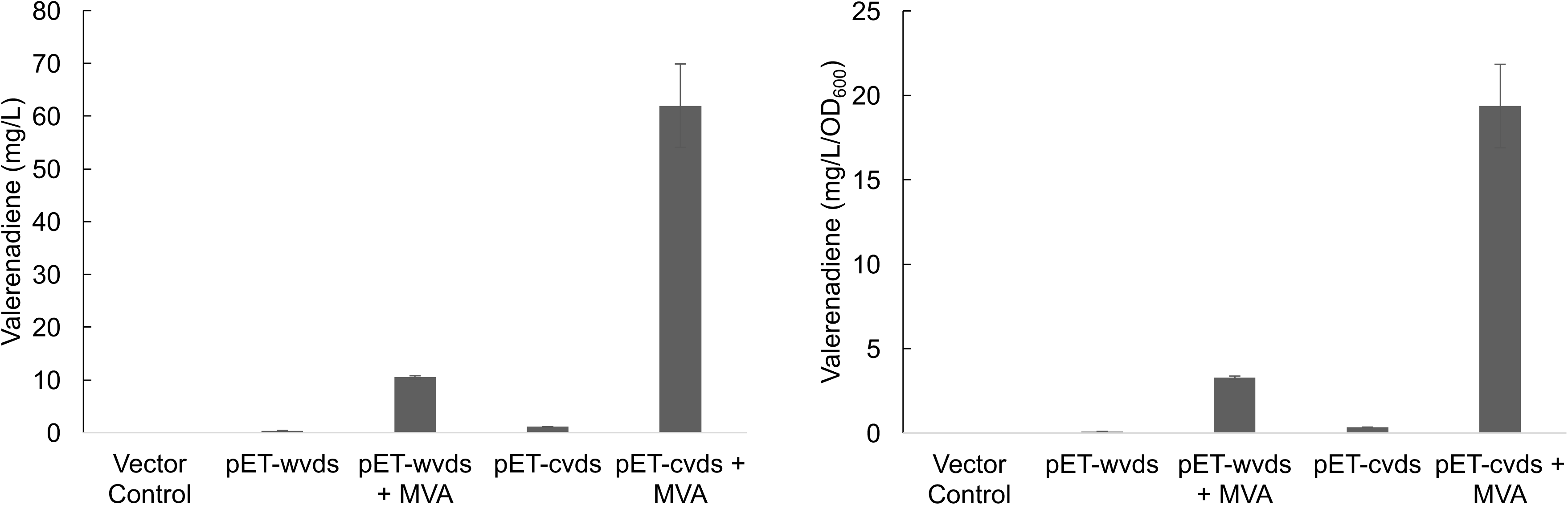
Production of valerenadiene using the T7 RNA polymerase to drive expression of *vds*. *E. coli* BL21(DE3) strains harboring empty vector control, pET-*wvds*, and pET-*cvds* were expressed with and without the *Saccharomyces cerevisiae* MVA pathway (pBBa5c-MevT-MBIS(CO, IspA)). Strains were grown in triplicate 5 mL 2xYT media + 3% glycerol fermentations with 1 mL decane overlay for 48 hours. (Left panel) valerenadiene volumetric production (mg/L) was determined after 48 hours. (Right panel) Valerenadiene production was normalized by dividing volumetric production by the biomass after 48 hours (mg/L/OD_600_).

## Discussion

Valerenic acid is the most potent sesquiterpenoid component produced in the roots of *Valeriana officinalis* that exhibits GABA-A activity. Development of this compound as a novel anxiolytic drug is hindered by the low production of valerenic acid by the native plant. As a first step towards the development of a production platform for valerenic acid analogues, we have employed *E. coli* as a workhorse metabolic host for engineering of terpene chemicals. In this study, valerenadiene was produced by introducing a codon-optimized valerenadiene synthase gene into *Escherichia coli* along with an engineered MEP pathway or *Saccharomyces cerevisiae*-based mevalonic acid pathway. The codon-optimized valerenadiene synthase gene was found to be a critical component for engineering high valerenadiene titers, as it resulted in 3-fold higher production of valerenadiene as compared to the wildtype sequence. This very likely indicates that translation of the codon-optimized form of the gene is very efficient. Further improvements in productivity were realized by the overexpression of the *dxs*, *idi*, and *fps* genes to augment carbon flux through the MEP pathway. This strategy resulted in a 65-fold increase in production of valerenadiene (2.09 mg/L). This was improved a further 6-fold by optimizing the fermentation medium to 2xYT with 3% glycerol supplementation (11.0 mg/L). Next, the *cvds* gene was fused to the stronger P_trc_ promoter, which resulted in approximately 40% increase in production of valerenadiene when pTrcHis-*cvds* was co-expressed with the engineered MEP pathway (20 mg/L). This result demonstrated the importance of driving expression of valerenadiene synthase from strong promoters to ensure ample availability of terpene synthase catalyst.

Notably, the valerenadiene synthase exhibits similar kinetic properties (e.g. K_m_ for FPP = 7.2 μM, *k*_cat_ = 5.7 x 10^-3^ s^-1^) to other published terpene synthases (Yeo et al., 2013). For example, the well-characterized amorphadiene synthase exhibits a two-fold lower K_m_ for FPP (i.e. K_m_ = 3.3 μM and *k*_cat_ = 6.8 x 10^-3^ s^-1^) (Picaud et al., 2005). Further efforts to improve the catalytic functionality of valerenadiene synthase could incorporate rationale modifications to the enzyme. Alternatively, further improvements in terpene synthase activity have been realized via expression of a terpene synthase as a gene fusion with farnesyl pyrophosphate synthase to achieve substrate channeling of FPP to the active site of VDS. Wang and co-workers reported that expression of *ispA* and apple farnesene synthase (*aFS*) from the same construct with the mevalonic acid pathway resulted in production of 57.6 mg/L of α-farnesene in *E. coli* (Wang et al., 2011). Fusing of *ispA*-*aFS* as a single gene construct with a (GGGGS)2 amino acid linker increased production to 87.8 mg/L of α-farnesene. Additionally, Niehaus et al. achieved substrate channeling by fusing the triterpene synthases SSL-1 and SSL-3 genes to form a single gene fusion, SSL-1+SSL-3, which increased production from 20 mg/L botryococcene to 50 mg/L botryococcene in yeast (Niehaus et al., 2011).

However, the greatest increases in valerenadiene yield involved heterologous expression of the *Saccharomyces cerevisiae* mevalonic acid pathway. Expression of the MVA pathway resulted in an unregulated metabolic flux towards FPP, which could be efficiently coupled to valerenadiene synthase for production of 62.0 mg/L valerenadiene in culture tubes with a specific productivity of 19.4 mg/L/OD_600_. This study showed that *E. coli* is capable of high-level production of valerenadiene with relatively minimal optimization of the production media and commercially available expression vectors, such as pBBa5c-MevT-MBIS(CO,IspA). Further efforts to improve valerenadiene yield will require optimization of the upstream and downstream genetic components of the MVA pathway and subsequent scale-up fermentation in bioreactors. In addition, we are conducting rational attempts to engineer CYP450s to oxidize and functionalize the valerenadiene skeleton to generate a library of potential GABA-A ligands.

## Acknowledgements

The authors gratefully acknowledge Scott Kinison, Garrett Zinck, and Dr. Joseph Chappell (University of Kentucky) for the pET-*wvds* and cloning of the pET-*cvds* constructs, valerenadiene standard, and helpful discussions concerning GC-MS protocols.

